# Antibiotic resistant and extended-spectrum β-lactamase producing faecal coliforms in wastewater treatment plant effluent

**DOI:** 10.1101/838136

**Authors:** Cian Smyth, Aidan O’Flaherty, Fiona Walsh, Thi Thuy Do

## Abstract

Wastewater treatment plants (WWTPs) provide optimal conditions for the maintenance and spread of antibiotic resistant bacteria (ARB) and antibiotic resistance genes (ARGs). In this work we describe the occurrence of antibiotic resistant faecal coliforms and their mechanisms of antibiotic resistance in the effluent of two urban WWTPs in Ireland. Effluent samples were collected from two WWTPs in Spring and Autumn of 2015 and 2016. The bacterial susceptibility patterns to 13 antibiotics were determined. The phenotypic tests were carried out to identify AmpC or extended-spectrum β-lactamase (ESBL) producers. The presence of ESBL genes were detected by PCR. Plasmids carrying ESBL genes were transformed into *Escherichia coli* DH5α recipient and underwent plasmid replicon typing to identify incompatibility groups. More than 90% of isolated faecal coliforms were resistant to amoxicillin and ampicillin, followed by tetracycline (up to 39.82%), ciprofloxacin (up to 31.42%) and trimethoprim (up to 37.61%). Faecal coliforms resistant to colistin and imipenem were detected in all effluent samples. Up to 53.98% of isolated faecal coliforms expressed a multi-drug resistance (MRD) phenotype. AmpC production was confirmed in 5.22% of isolates. The ESBL genes were confirmed for 11.84% of isolates (9.2% of isolates carried *bla*_TEM_, 1.4% *bla*_SHV-12_, 0.2% *bla*_CTX-M-1_ and 1% *bla*_CTX-M-15_). Plasmids extracted from 52 ESBL isolates were successfully transformed into recipient *E. coli*. The detected plasmid incompatibility groups included the IncF group, IncI1, IncHI1/2 and IncA/C. These results provide evidence that treated wastewater is polluted with ARB and MDR faecal coliforms and are sources of ESBL-producing, carbapenem and colistin resistant *Enterobacteriaceae*.

**Importance:** Antibiotic resistant bacteria (ARB) are an emerging environmental concern with a potential impact on human health. The results provide the evidence that treated wastewater is polluted with antibiotic resistant bacteria containing mobile resistance mechanisms of importance to clinical treatment of pathogens and multi-drug resistant (MDR) faecal coliforms. They are sources of relatively high proportions of ESBL-producing *Enterobacteriaceae*, and include carbapenem and colistin resistant *Enterobacteriaceae.* The significance of this study is the identification of the role of WWTPs as a potential control point to reduce or stop the movement of ESBL, MDR and colistin resistant bacteria into the environment from further upstream sources, such as human or animal waste.

## Introduction

The dissemination of antimicrobial resistance within bacterial communities and the selection of new resistance mechanisms are due to the large-scale use of antibiotics in agricultural, veterinary and human clinical applications (1-8). The emergence of antibiotic resistant bacteria (ARB) is a major public health issue which poses a serious therapeutic challenge worldwide (6). Wastewater treatment plants (WWTPs) are considered potential sources of ARB and antibiotic resistance genes (ARGs), which play an important role in the spread of antibiotic resistance into the environment (9, 10). The urban WWTPs receive wastewater from human communities, which contains high concentrations of chemical matter, including antibiotics and microorganisms, including ARB. Therefore, WWTPs are favourable environments with optimal conditions for the development and spread of ARB and ARGs (11, 12). Both ARB and ARGs were detected in wastewater samples globally (13-24). However, little is known of the fate of these bacteria; and the role of WWTPs in releasing ABR and ARGs into the environment through treated effluent (13). A recent report by Flach et al shows no evidence for the selection of antibiotic resistance in WWTPs (25); however, large amounts of resistant bacteria were identified throughout the wastewater treatment process (7, 26). The conventional wastewater treatment process can remove some ARB (27, 28), but ARB were still found in large proportions in the effluent (18, 29). In some cases, ARB were detected at higher rates in WWTP effluent than in the influent (30, 31).

AmpC cephalosporinases and extended-spectrum β-lactamases (ESBLs) are some of the most clinically important antibiotic resistance mechanisms (32). The dissemination of AmpC or ESBL producing *Escherichia coli* were identified in different types of aquatic environments, particularly in wastewater (33-36). The prevalence of AmpC or ESBL producing bacteria pose a global health problem due to limitations of therapeutic options for the treatment of infections caused by these bacteria (37). The ESBL genes are frequently located on mobile genetic elements (38). Among more than 300 subtypes of ESBL genes, *bla*_TEM_ and *bla*_SHV_ groups were the most common ESBL genes identified in human pathogens until the late 1990s. These groups were replaced by *bla*_CTX-M_ genes since the beginning of the 2000s and *Escherichia coli* became the most prevalent ESBL producing bacteria among clinical *Enterobacteriaceae* (39). The monitoring of antibiotic resistance from WWTPs provides the information required to track the dissemination of ARB and ARGs into the environment (40, 41). Moreover, the analysis of ARB and ARGs in urban WWTPs is considered as an alternative method for the indirect study of antibiotic resistance in human populations from which the WWTPs receive wastewater (42). Indeed, the resistance rates of indicator bacteria in wastewater may give useful information to identify the changes in resistance in the human populations (43). The objectives of this study were to characterise the faecal coliforms resistome leaving urban WWTPs via the effluent. This was achieved through i) assessment of the prevalence of antibiotic resistant faecal coliforms in the effluent from two Irish urban WWTPs, ii) characterisation of the antibiotic resistance profiles of these bacteria, iii) identification of the occurrence of AmpC or ESBL producing faecal coliforms and iv) identification of the resistance mechanisms and their potential mobility.

## Materials and Methods

### Isolation of total faecal coliforms

Final effluent samples were collected from two urban WWTPs in Ireland during early Spring and late Autumn in 2015 and 2016. These WWTPs were representative of medium sized WWTPs with 100% urban agglomerations, include tertiary treatment, and the distance between them was less than 100km. Faecal coliforms were isolated using the membrane filtration method (44). The effluent samples (1 mL and 10 mL) were filtered through nitrocellulose membranes (Sigma Aldrich/Merck). The filters were then incubated on mFC agar at 37 °C for 24 hours in the presence or absence of antibiotics: amoxicillin (32 mg/L), ciprofloxacin (1 mg/L) or tetracycline (16 mg/L). All procedures were performed in triplicate.

### Antibiotic susceptibility test using agar dilution and disk diffusion methods

Bacterial isolates were subjected to antibiotic susceptibility testing using the agar dilution method following the CLSI recommendations (45). The antibiotics tested were tetracycline, amoxicillin, ampicillin, ciprofloxacin, kanamycin, gentamicin, colistin, chloramphenicol and trimethoprim. The imipenem, meropenem, cefotaxime and ceftazidime susceptibilities were determined using the disk diffusion method (45). The minimum inhibitory concentration (MIC) breakpoints for Enterobacteriaceae in the CLSI guidelines (45) were used to identify ARB. The EUCAST MIC breakpoint for colistin was used (46). Bacterial isolates resistant to three or more different antibiotic classes of antibiotics were defined as multidrug resistant. The resistance percentages of bacteria were calculated as: percentage (%) = [(Number of resistant faecal coliforms to an antibiotic/ total number of tested faecal coliforms) X 100].

### Phenotypic identification of the production of Metallo-beta-lactamase (MBL), ESBL and AmpC enzymes

Bacteria resistant to imipenem and/or meropenem were subjected to the Imipenem-EDTA double-disk synergy test as described previously (47). Isolates resistant to cefotaxime and/or ceftazidime were subjected to ESBL testing following the CLSI guidelines and AmpC testing using phenylboronic acid (45, 48).

### Identification of antibiotic resistance genes and bacterial species

Putative MBL producing carbapenem resistant isolates (identified using the imipenem-EDTA double-disk synergy test) were subjected to multiplex PCR to identify the carbapenem resistance genes. The primer sets included bla_GES_, bla_GIM_, bla_IMI_, bla_IMP_, bla_KPC_, bla_NDM_, bla_OXA-23_, bla_OXA-40_, bla_OXA-48_, bla_OXA-51_, bla_OXA-58_, bla_VIM_ (Table 1) (49). Isolates displaying a positive ESBL phenotype and were phenotypically negative for AmpC production were further analysed using the ESBL multiplex-PCR. The primer sets were used to detect the *bla*_TEM_, *bla*_SHV_, and *bla*_CTX-M-_group genes 1, 2, 8, 9, and 25 (Table 1) (50, 51).

**Table 1:**
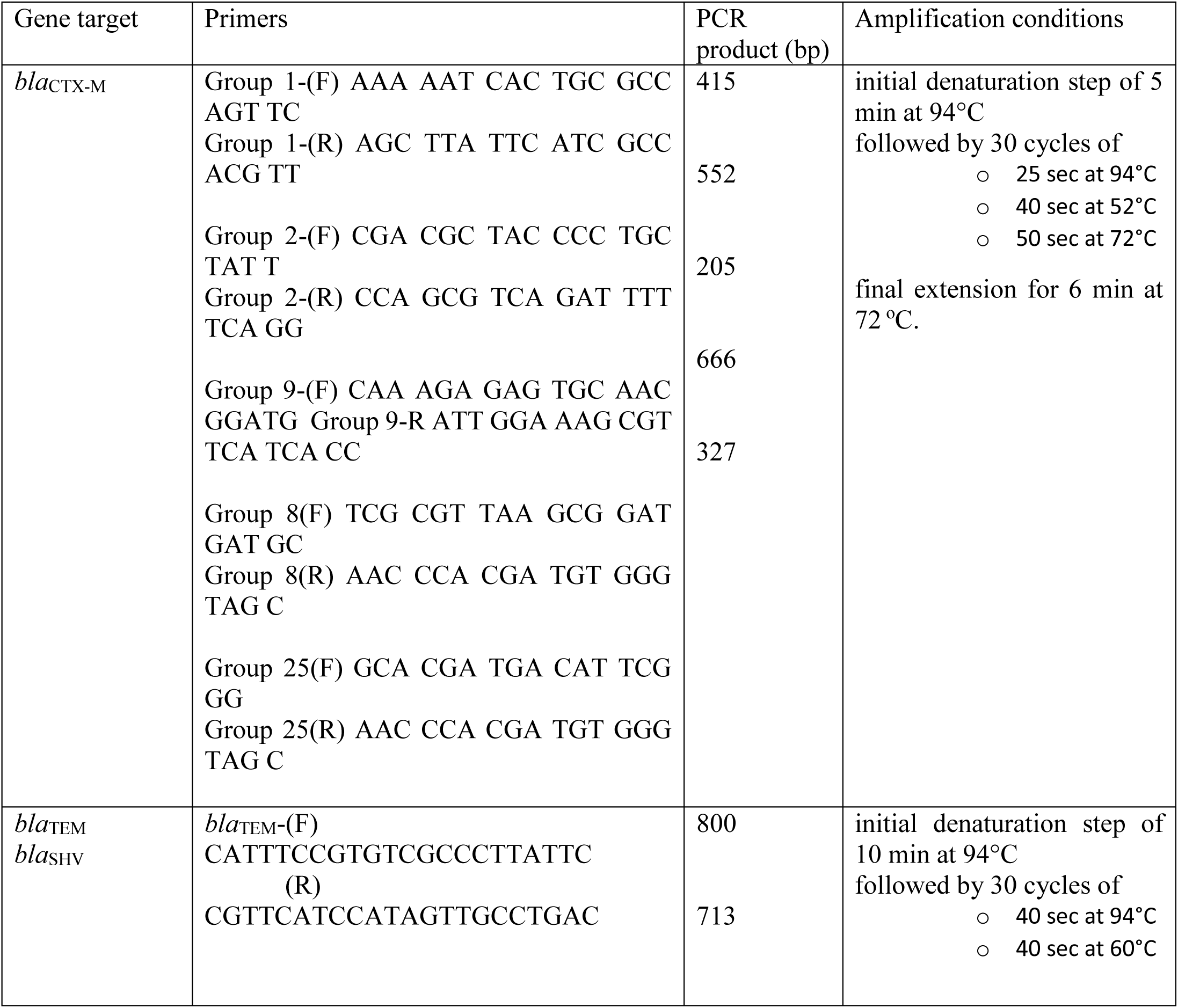
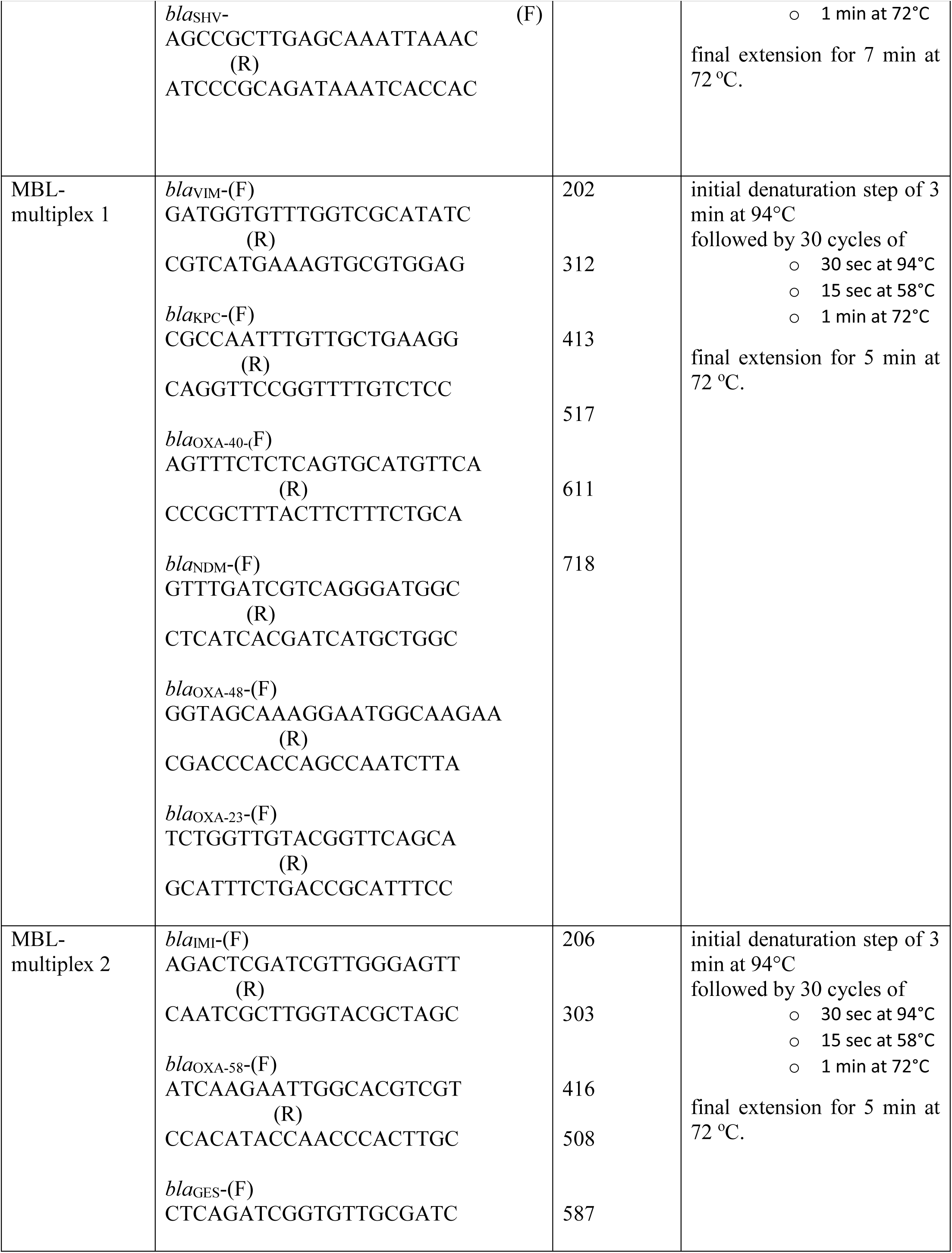
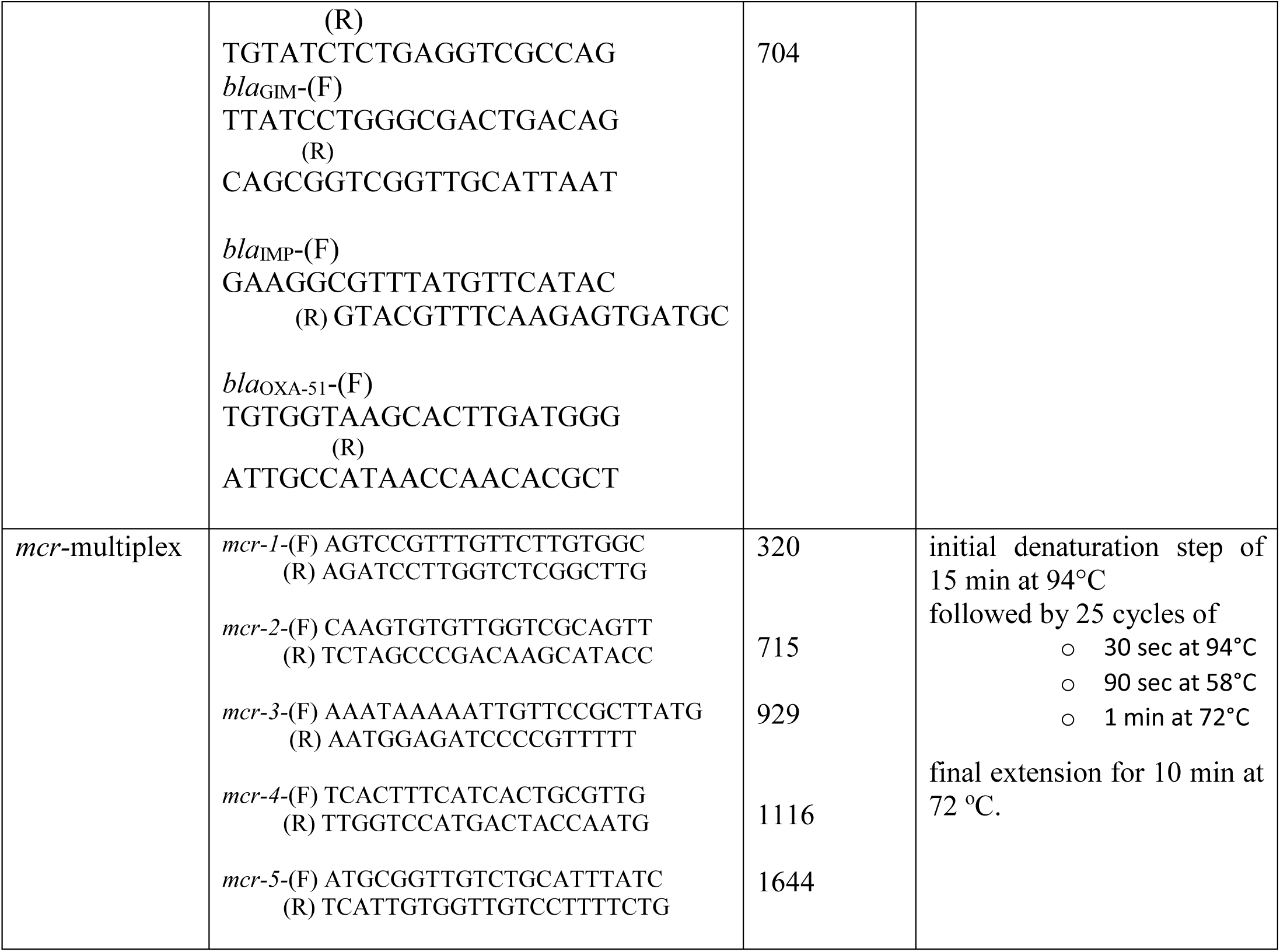
Primers used in multiplex PCRs to detect ESBL (50, 51), MBL (49) and *mcr*-genes (52)

Bacterial isolates resistant to colistin were further analysed by for the presence of the plasmid mediated colistin resistance mechanisms. Five primer sets were used to screen for the presence of *mcr*-1, 2, 3, 4 and 5, as recommended by the EU reference laboratory-antimicrobial resistance (Table 1) (52). The bacterial species of isolates carrying ESBL genes was identified by Sanger sequencing of the V3 and V4 region of bacterial 16S rRNA using forward primer: 5’-TCGTCGGCAGCGTCAGATGTGTATAAGAGACAGCCTACGGGNGGCWGCAG-3’ and reverse primer: 5’-GTCTCGTGGGCTCGGAGATGTGTATAAGAGACAGGACTACHVGGGTATCTAATC C-3’ (53).

### Plasmid transfer by transformation and replicon typing

Plasmids were extracted from ESBL positive isolates carrying the *bla*_TEM_, *bla*_SHV_, and *bla*_CTX-M_ genes using the Macherey-Nagel NucleoSpin plasmid isolation kit. The plasmids were transformed into *Escherichia coli* DH5α using heat-shock transformation (54, 55). The transformants were selected on LB agar supplemented with ampicillin (32 mg/L). The presence of *bla*_TEM_, *bla*_SHV_, and *bla*_CTX-M_ genes in transformants were confirmed by PCR. All transformants were subjected to antimicrobial susceptibility testing against imipenem, meropenem, ertapenem, ciprofloxacin, chloramphenicol, tetracycline, amikacin, gentamycin, kanamycin, trimethoprim and colistin. Replicon typing via PCR were performed on ESBL transformants with 18 pairs of primers recognizing FIA, FIB, FIC, HI1, HI2, I1-Iγ, L/M, N, P, W, T, A/C, K, B/O, X, Y, F, and FIIA in 3 multiplex panels (56).

## Results

### Antibiotic susceptibility patterns

In total, 498 faecal coliforms were isolated from all WWTP effluent samples, comprising 226 isolates from WWTP A and 272 from WWTP B. These isolates were subjected to antibiotic susceptibility testing. Among the tested β-lactam antibiotics, more than 90% of bacteria isolated from the two WWTPs were resistant to amoxicillin and ampicillin (Table 2, Figure 1); greater than 20% were resistant to cefotaxime or ceftazidime. All ceftazidime resistant isolates were also resistant to cefotaxime. Carbapenem resistance was detected at relatively lower levels (Table 2). Colistin resistance was found at a higher percentage in WWTP B effluent than in WWTP A. We also identified that there were no differences in the identification of colistin resistant isolates in antibiotic susceptibility testing by the agar dilution method compared to the microbroth dilution method. Multi-drug resistant (MDR) faecal coliforms were detected at approximately 50 % of the total isolates tested (Table 2). The resistance prevalence to other antibiotics were found at similar levels between the two WWTPs.

**Table 2:**
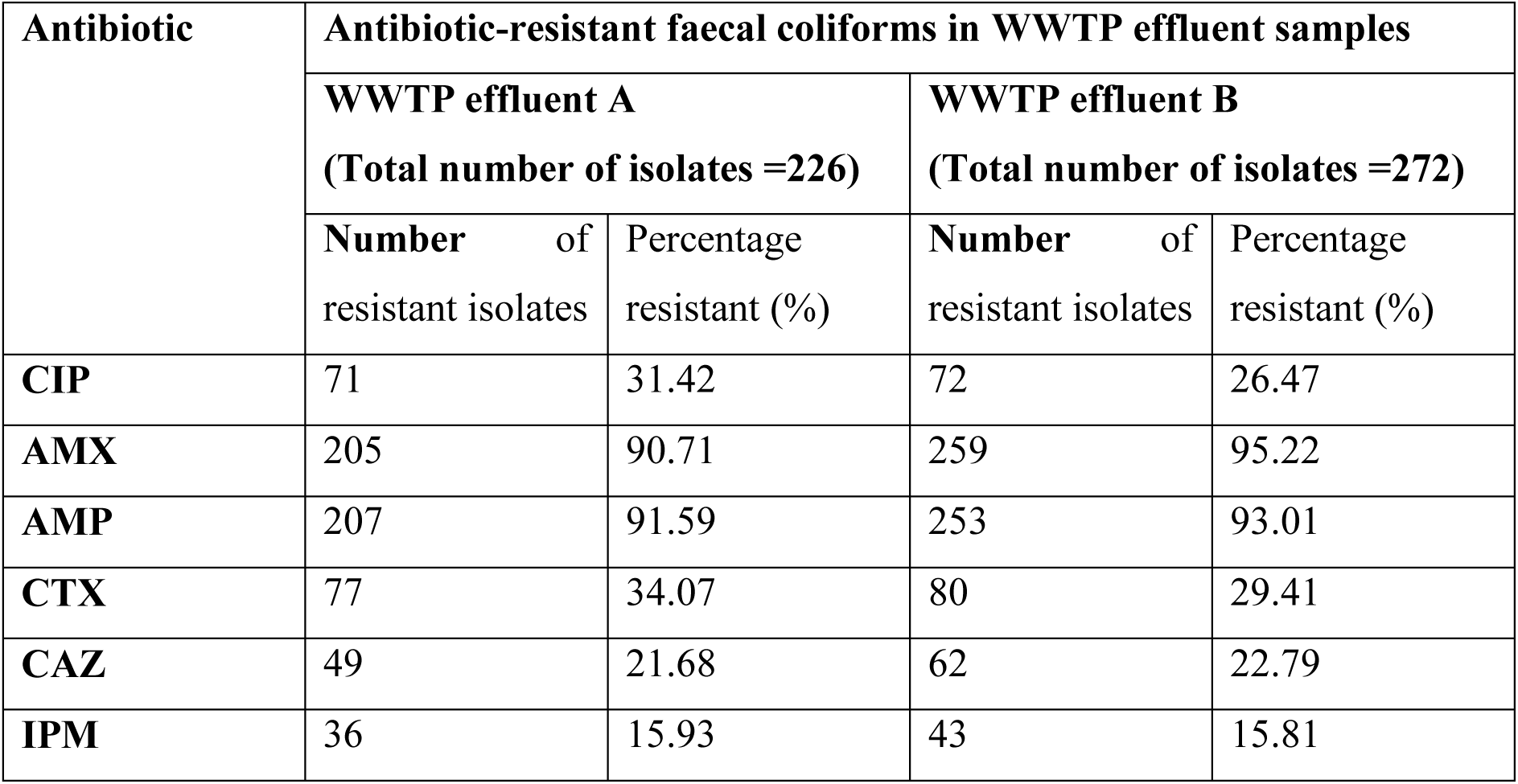
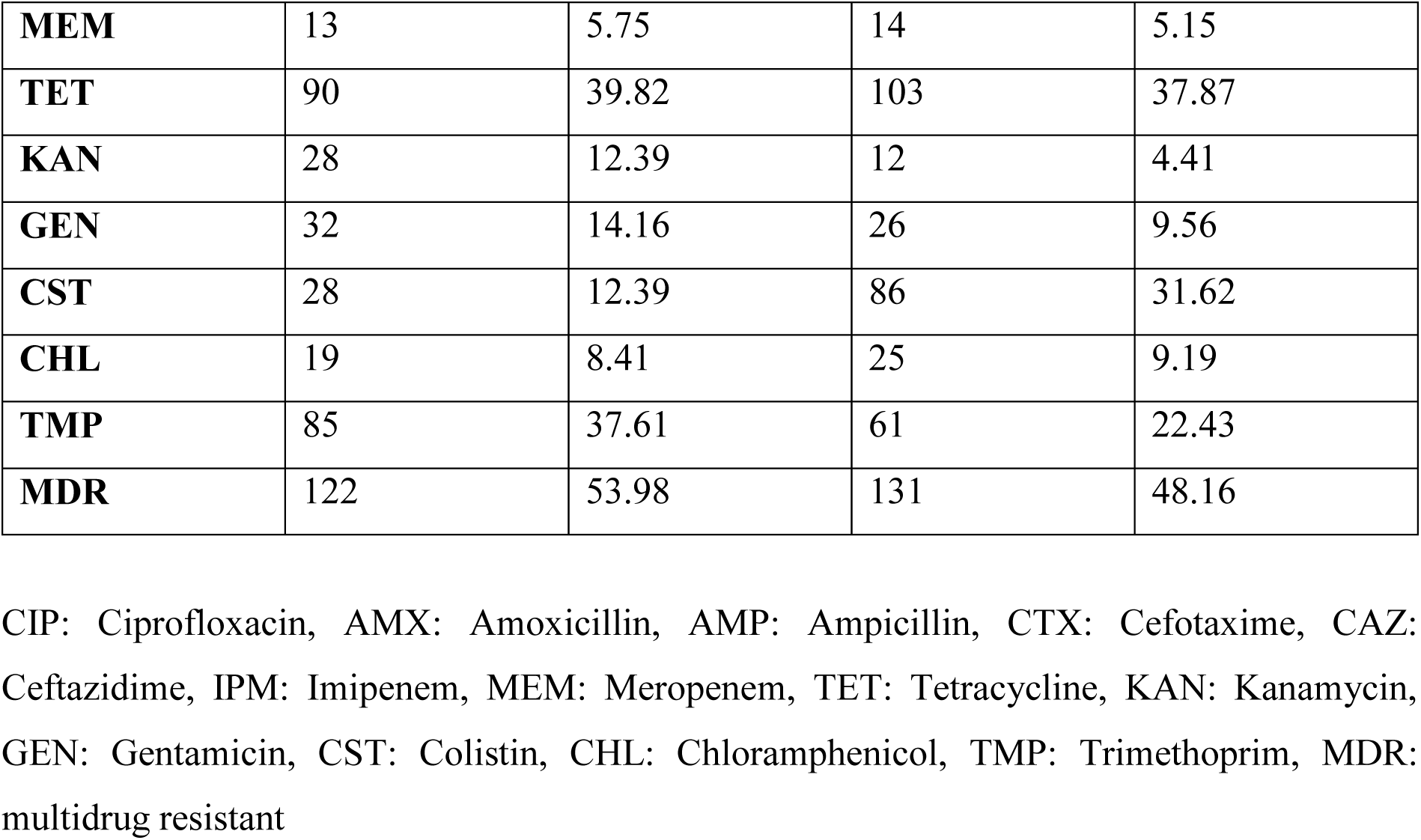
Prevalence of antibiotic resistance phenotypes in faecal coliforms isolated from WWTP effluent samples

**Figure 1:**
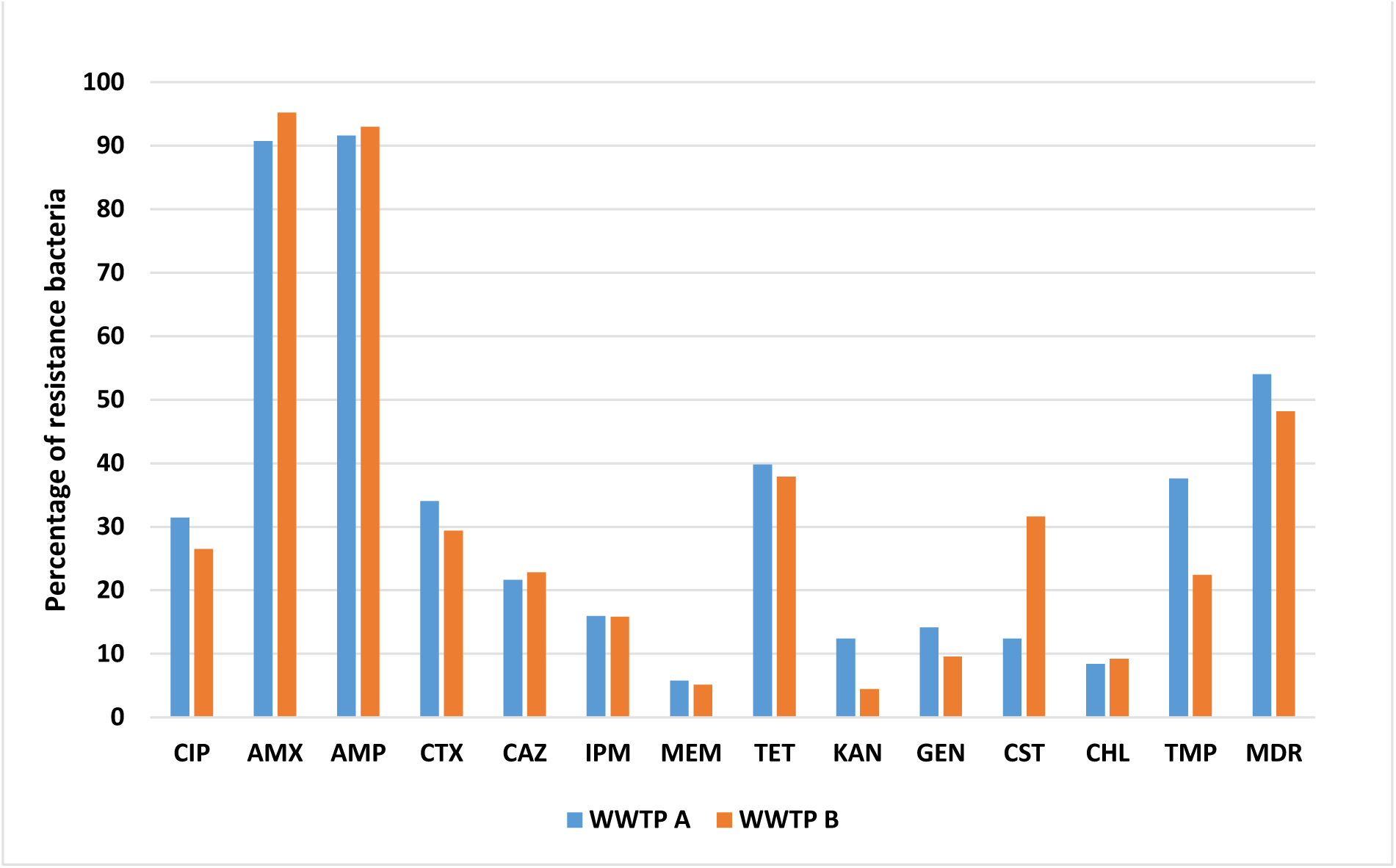
Percentage of faecal coliforms isolated from WWTP effluent samples identified with antibiotic resistance phenotypes. CIP: Ciprofloxacin, AMX: Amoxicillin, AMP: Ampicillin, CTX: Cefotaxime, CAZ: Ceftazidime, IPM: Imipenem, MEM: Meropenem, TET: Tetracycline, KAN: Kanamycin, GEN: Gentamicin, CST: Colistin, CHL: Chloramphenicol, TMP: Trimethoprim, MDR: multidrug resistant

### Detection of antibiotic resistance genes and speciation

Faecal coliforms with resistance to cefotaxime and ceftazidime were considered putative ESBL-producers (n = 157). AmpC production was confirmed in 26 of these isolates (13 from WWTP A and 13 from WWTP B). Of the 131 remaining isolates, 89 (39 from WWTP A and 50 from WWTP B) were identified as phenotypic ESBL producers. The ESBL multiplex PCR revealed the presence of *bla*_TEM_, *bla*_SHV-12_ and *bla*_CTX-M_ group 1 genes in 62 isolates. Among them, 49 isolates carried *bla*_TEM_ (25 from WWTP A and 24 from WWTP B), 7 *bla*_SHV-12_ (3 from WWTP A and 4 from WWTP B), 1 *bla*_CTX-M-1_ (WWTP B) and 5 *bla*_CTX-M-15_ (2 from WWTP A and 3 from WWTP B) (Table 3). Almost all ESBL producers were *E. coli* (46 carrying *bla*_TEM_, 2 carrying *bla*_SHV-12,_ 1 carrying *bla*_CTX-M-1_ and 2 carrying *bla*_CTX-M-15_). In addition, 7 isolates were *Klebsiella spp*. (5 carrying *bla*_SHV_, 2 carrying *bla*_CTX-M-15_) and 1 *bla*_CTX-M-15_ positive *Enterobacter spp*.

**Table 3:** Extended-spectrum β- lactamase genes identified in faecal coliforms isolated from WWTP effluent samples

| **ESBL gene** | **WWTP A**<br>**Number of ESBL**<br>**isolates** | **WWTP B**<br>**Number of ESBL**<br>**isolates** |
| --- | --- | --- |
| *bla*<sub>TEM-116</sub> | 6 | 5 |
| *bla*<sub>TEM-12</sub> | 3 | 2 |
| *bla*<sub>TEM-1-like</sub> (>99% identical to *bla*<sub>TEM-143</sub> or *bla*<sub>TEM-164</sub>) | 16 | 17 |
| *bla*<sub>CTX-M-1</sub> | 0 | 1 |
| *bla*<sub>CTX-M-15</sub> | 2 | 3 |
| *bla*<sub>SHV-12</sub> | 3 | 4 |

In total, 79 isolates (36 from WWTP A and 43 from WWTP B) resistant to imipenem were screened for MBL production. Metallo-beta-lactamase production was identified in 36 isolates. However, all isolates were negative for known MBL genes, using the MBL multiplex PCR. More than 100 colistin resistant isolates were detected. However, none of these isolates were positive for the *mcr*-genes using the *mcr*-targeted PCR analysis.

### Plasmid transformation, plasmid resistance profile and replicon typing

The plasmids were extracted from 62 ESBL faecal coliform isolates (30 from WWTP A and 32 from WWTP B). Plasmids extracted from 52 ESBL isolates (46 plasmids carried *bla*_TEM_, 2 plasmids *bla*_CTX-M15_, 1 *bla*_CTX-M-1_ and 3 with *bla*_SHV-12_) were successfully transferred into *E. coli* Dh5α recipients. The plasmids extracted from the 10 ESBL isolates (3 with *bla*_TEM_, 3 with *bla*_CTX-M15_ and 4 with *bla*_SHV-12_) could not be transferred, suggesting that the ESBL genes identified in these isolates are located on the bacterial chromosomes.

All transformants were sensitive to the tested carbapenem antibiotics (imipenem, meropenem and ertapenem) and gentamicin, 16 were resistant to chloramphenicol, 16 resistant to tetracycline, 10 resistant to trimethoprim, 3 resistant to colistin, 2 resistant to kanamycin, 1 resistant to amikacin and 1 resistant to ciprofloxacin. Nine transformants show a resistance phenotype to two drugs from different classes and eight show a multidrug resistance phenotype. The plasmids from 21 transformants could be typed using PCR replicon typing. Of these, 11 plasmids were IncF replicon type (3 IncFIA, 1 IncFIIA, 1 IncFIB and 6 non-specific IncF group), 5 were IncI1 replicon type, 2 were IncHI1 replicon type, 2 were IncHI2 replicon type and 1 was IncA/C replicon type. The IncF group of replicons was the most prevalent across plasmids from both WWTPs. The IncI1 type was detected only in bacteria isolated from the WWTP B effluent samples.

## Discussion

Our work presents the antibiotic resistance patterns of faecal coliforms in the effluent of two WWTPs in Ireland. Wastewaters with faecal contamination are considered reservoirs for ARB and ARGs in the environment (57, 58). Among all tested antibiotics, resistance to amoxicillin and ampicillin was most prevalent, followed by tetracycline and, cefotaxime and ciprofloxacin. High levels of β-lactam resistance were previously detected in Enterobacteriaceae in an urban WWTP (59), and resistance to tetracycline and fluoroquinolones was found at lower rates. The higher resistance rate of *E. coli* to ampicillin and tetracycline as well as lower rates of resistance to ciprofloxacin were detected in WWTPs in Portugal (60). A study of raw sewage in Brazil identified 100% sensitivity of *E. coli* to ciprofloxacin and amoxicillin and tetracycline resistance levels in the range of 50 – 75% (61). The proportions of ciprofloxacin resistant faecal coliforms were 31.42% in WWTP A effluent and 26.47% in WWTP B. This resistance rate is higher in comparison to reported levels of ciprofloxacin resistance in the *E. coli* isolated from other WWTPs (59, 62, 63). In general, differences in antibiotic resistance percentages were observed between the two WWTPs, particularly for colistin, trimethoprim and kanamycin (Figure 1). As these two WWTPs are using the same treatment process, this difference may be associated with their location or other external factors.

The presence of colistin resistant and carbapenem resistant isolates in tested WWTP effluents raises the possibility of transferability or risk to human health. These antibiotics are known as ‘last resort’ antibiotics to treat MDR bacteria. Previous studies on colistin resistant Enterobacteriaceae focus mainly on food, human and animal samples (64-67). To date, there are only a few studies conducted on the identification of *mcr* genes in waterborne bacteria (68-72). 114 faecal coliform isolates resistant to colistin (28 from WWTP A and 86 from WWTP B) were detected in our work, none of them were positive for the *mcr* genes. Among colistin resistant isolates, the plasmids extracted from 43 isolates (4 from WWTP A and 39 from WWTP B) were successfully transferred to the recipient *E. coli* DH5α in the transformation studies. The proportion of colistin resistant coliforms in our study was lower than those in the studies of Igwaran et al.(68), where they found approximately 60% of isolates with resistance to colistin. Carbapenem resistant Enterobacteriaceae were studied in hospital wastewater and in WWTPs previously (73-76). The percentage of carbapenem resistance phenotypes in our work were considered high in comparison to previous studies in wastewater (33, 77). However, there were no known carbapenem resistance genes detected in the carbapenem resistant coliforms. This suggests that other resistance mechanisms are responsible for the resistance phenotypes and thus require further study to characterise these novel mechanisms.

Multi-drug resistant faecal coliforms were retrieved at high rates from all effluent samples. In the study of Lefkowitz and Durán (78), 60% of *E. coli* in WWTP effluent were resistant to two or more antibiotics and 25% to four or more antibiotics. The study of Garcia *et al* (79) in WWTP effluent showed no more than 12% of *E. coli* were resistant to two antibiotics and less than 10% to three or more antibiotics. *Escherichia coli* (34.3%) were found to be resistant to two or more antibiotics and 8.8% to four or more antibiotics in treated wastewater in Portugal (59). The MDR faecal coliforms in our study were found in the same range of the study of Lefkowitz and Durán, but at a higher percentage than in others.

The ESBLor AmpC producing faecal coliforms were recovered from all WWTP effluent samples. The rate of ESBL producing *Enterobacteriacea* in our work were within the range of previous studies. It was considerably high in comparison to some studies of WWTP effluent (0.5-9.8%) (34, 80-83). However it was lower than those studies in wastewater (45-100%) (84, 85). The high load of bacteria and rich nutrient environment in WWTPs facilitates the transfer of ARGs among bacteria (11, 12). These may explain the relatively high rates of ESBL producers in WWTP effluent.

The *bla*_TEM_ were the most prevalent beta-lactamase in this study, which is similar to a study on hospital wastewater in Brazil (85). However, this is in contrast with other findings with *bla*_CTX-M_ being the most frequent ESBL genes from various samples including in hospital effluent, surface water and WWTPs (34, 39, 86). This result was confirmed by qPCR data performed in another study in the framework of the StARE project which showed the detection of *bla*_TEM_ genes, where *bla*_SHV_ and *bla*_CTX-M_ group 1 were not detected (22). Most of ESBL genes were found in *E. coli*, others were carried by *Klebsiella spp.* in our work. These results are in an agreement with previous findings which indicated that *E. coli* were the most common ESBL-producers among Enterobacteriaceae (39).

Transformation of plasmids carrying ESBL genes were successful for 88% of ESBL donor isolates in this work. Different replicon variants were found in the ESBL plasmids. The most prevalent replicon was IncF group which were also reported in other studies of plasmid replicon typing in Enterobacteriaceae (87-89). These replicons have a narrow host range and can be transfer easily among *E. coli* species (90). The cross-resistance of ESBL producers to other antibiotic causes of great concern as ESBL genes are frequently located on conjugative plasmids carrying other ARGs (91). In this work eight of 52 transferable plasmids carrying ESBL genes expressed a MDR phenotype.

## Conclusion

Effluent samples from two WWTPs demonstrated the presence of ARB and MDR and, of particular importance, a source of a relatively high proportion of ESBL-producing, carbapenem and colistin resistant Enterobacteriaceae. Although the bacteria were phenotypically resistant to colistin or carbapenems no known mobile resistance mechanisms were detected, despite the ability to transfer the resistance phenotype via transformation. Thus, faecal coliforms from WWTP effluent are sources of novel ARGs conferring resistance to antibiotics of last line of defence. The ability of these bacteria to survive in water has been demonstrated for many years. The significance of this study is the identification of the role of WWTPs as a potential control point to reduce or stop the movement of resistant bacteria and genes into the environment from further upstream sources, such as human or animal waste. In addition, this enables the use of additional treatment technologies to be added to WWTPs to stop or reduce such ARB and ARGs entering the water environments globally.

## Acknowledgments

This work was funded by the Irish Environmental Protection Agency, in the frame of the collaborative international consortium of Water challenges for a changing world Joint Programming Initiative (Water JPI) Pilot Call, project Stare and the EPA – Maynooth University co-fund project “Survival of mobile antibiotic resistance in water”. COST Action ES1403: New and emerging challenges and opportunities in wastewater reuse (NEREUS).

## References

1. Alonso A, Sanchez P, Martinez JL. 2001. Environmental selection of antibiotic resistance genes. Environ Microbiol 3:1–9.

2. Larson E. 2007. Community factors in the development of antibiotic resistance. Annu Rev Public Health 28:435–47.

3. Aminov RI. 2009. The role of antibiotics and antibiotic resistance in nature. Environ Microbiol 11:2970–88.

4. Taylor NG, Verner-Jeffreys DW, Baker-Austin C. 2011. Aquatic systems: maintaining, mixing and mobilising antimicrobial resistance? Trends Ecol Evol 26:278–84.

5. Udikovic-Kolic N, Wichmann F, Broderick NA, Handelsman J. 2014. Bloom of resident antibiotic-resistant bacteria in soil following manure fertilization. Proc Natl Acad Sci U S A 111:15202–7.

6. Berendonk TU, Manaia CM, Merlin C, Fatta-Kassinos D, Cytryn E, Walsh F, Burgmann H, Sorum H, Norstrom M, Pons MN, Kreuzinger N, Huovinen P, Stefani S, Schwartz T, Kisand V, Baquero F, Martinez JL. 2015. Tackling antibiotic resistance: the environmental framework. Nat Rev Microbiol 13:310–7.

7. Rizzo L, Manaia C, Merlin C, Schwartz T, Dagot C, Ploy MC, Michael I, Fatta-Kassinos D. 2013. Urban wastewater treatment plants as hotspots for antibiotic resistant bacteria and genes spread into the environment: a review. Sci Total Environ 447:345–60.

8. Sharma VK, Johnson N, Cizmas L, McDonald TJ, Kim H. 2016. A review of the influence of treatment strategies on antibiotic resistant bacteria and antibiotic resistance genes. Chemosphere 150:702–714.

9. Baquero F, Martinez JL, Canton R. 2008. Antibiotics and antibiotic resistance in water environments. Curr Opin Biotechnol 19:260–5.

10. Ferro G, Guarino F, Cicatelli A, Rizzo L. 2017. beta-lactams resistance gene quantification in an antibiotic resistant Escherichia coli water suspension treated by advanced oxidation with UV/H2O2. J Hazard Mater 323:426–433.

11. Hultman J, Tamminen M, Parnanen K, Cairns J, Karkman A, Virta M. 2018. Host range of antibiotic resistance genes in wastewater treatment plant influent and effluent. FEMS Microbiol Ecol 94.

12. Ham YS, Kobori H, Kang JH, Matsuzaki T, Iino M, Nomura H. 2012. Distribution of antibiotic resistance in urban watershed in Japan. Environ Pollut 162:98–103.

13. Bouki C, Venieri D, Diamadopoulos E. 2013. Detection and fate of antibiotic resistant bacteria in wastewater treatment plants: a review. Ecotoxicol Environ Saf 91:1–9.

14. Goni-Urriza M, Pineau L, Capdepuy M, Roques C, Caumette P, Quentin C. 2000. Antimicrobial resistance of mesophilic Aeromonas spp. isolated from two European rivers. J Antimicrob Chemother 46:297–301.

15. Kummerer K. 2009. Antibiotics in the aquatic environment--a review--part II. Chemosphere 75:435–41.

16. Munir M, Wong K, Xagoraraki I. 2011. Release of antibiotic resistant bacteria and genes in the effluent and biosolids of five wastewater utilities in Michigan. Water Res 45:681–93.

17. Novo A, Andre S, Viana P, Nunes OC, Manaia CM. 2013. Antibiotic resistance, antimicrobial residues and bacterial community composition in urban wastewater. Water Res 47:1875–87.

18. Pruden A, Pei R, Storteboom H, Carlson KH. 2006. Antibiotic resistance genes as emerging contaminants: studies in northern Colorado. Environ Sci Technol 40:7445–50.

19. Schwartz T, Kohnen W, Jansen B, Obst U. 2003. Detection of antibiotic-resistant bacteria and their resistance genes in wastewater, surface water, and drinking water biofilms. FEMS Microbiol Ecol 43:325–35.

20. Cai L, Ju F, Zhang T. 2014. Tracking human sewage microbiome in a municipal wastewater treatment plant. Appl Microbiol Biotechnol 98:3317–26.

21. Thi Thuy Do Sinéad Murphy FW. 2017. Antibiotic Resistance and Wastewater Treatment Process – Do – – Wiley Online Books – Wiley Online Library. In Fugère PLKR (ed), Antimicrobial Resistance in Wastewater Treatment Processes. Wiley Blackwell.

22. Parnanen KMM, Narciso-da-Rocha C, Kneis D, Berendonk TU, Cacace D, Do TT, Elpers C, Fatta-Kassinos D, Henriques I, Jaeger T, Karkman A, Martinez JL, Michael SG, Michael-Kordatou I, O’Sullivan K, Rodriguez-Mozaz S, Schwartz T, Sheng H, Sorum H, Stedtfeld RD, Tiedje JM, Giustina SVD, Walsh F, Vaz-Moreira I, Virta M, Manaia CM. 2019. Antibiotic resistance in European wastewater treatment plants mirrors the pattern of clinical antibiotic resistance prevalence. Sci Adv 5:eaau9124.

23. Do TT, Delaney S, Walsh F. 2019. 16S rRNA gene based bacterial community structure of wastewater treatment plant effluents. FEMS Microbiol Lett 366.

24. Karkman A, Do TT, Walsh F, Virta MPJ. 2018. Antibiotic-Resistance Genes in Waste Water. Trends Microbiol 26:220–228.

25. Flach CF, Genheden M, Fick J, Joakim Larsson DG. 2018. A Comprehensive Screening of Escherichia coli Isolates from Scandinavia’s Largest Sewage Treatment Plant Indicates No Selection for Antibiotic Resistance. Environ Sci Technol 52:11419–11428.

26. Luczkiewicz A, Jankowska K, Fudala-Ksiazek S, Olanczuk-Neyman K. 2010. Antimicrobial resistance of fecal indicators in municipal wastewater treatment plant. Water Res 44:5089–97.

27. Guardabassi L, Lo Fo Wong DM, Dalsgaard A. 2002. The effects of tertiary wastewater treatment on the prevalence of antimicrobial resistant bacteria. Water Res 36:1955–64.

28. Martins da Costa P, Vaz-Pires P, Bernardo F. 2006. Antimicrobial resistance in Enterococcus spp. isolated in inflow, effluent and sludge from municipal sewage water treatment plants. Water Res 40:1735–40.

29. Hembach N, Schmid F, Alexander J, Hiller C, Rogall ET, Schwartz T. 2017. Occurrence of the mcr-1 Colistin Resistance Gene and other Clinically Relevant Antibiotic Resistance Genes in Microbial Populations at Different Municipal Wastewater Treatment Plants in Germany. Front Microbiol 8:1282.

30. Korzeniewska E, Harnisz M. 2018. Relationship between modification of activated sludge wastewater treatment and changes in antibiotic resistance of bacteria. Sci Total Environ 639:304–315.

31. Czekalski N, Berthold T, Caucci S, Egli A, Burgmann H. 2012. Increased levels of multiresistant bacteria and resistance genes after wastewater treatment and their dissemination into lake geneva, Switzerland. Front Microbiol 3:106.

32. Bonomo RA. 2017. beta-Lactamases: A Focus on Current Challenges. Cold Spring Harb Perspect Med 7.

33. Zurfluh K, Hachler H, Nuesch-Inderbinen M, Stephan R. 2013. Characteristics of extended-spectrum beta-lactamase- and carbapenemase-producing Enterobacteriaceae Isolates from rivers and lakes in Switzerland. Appl Environ Microbiol 79:3021–6.

34. Korzeniewska E, Harnisz M. 2013. Extended-spectrum beta-lactamase (ESBL)-positive Enterobacteriaceae in municipal sewage and their emission to the environment. J Environ Manage 128:904–11.

35. Diallo AA, Brugere H, Kerouredan M, Dupouy V, Toutain PL, Bousquet-Melou A, Oswald E, Bibbal D. 2013. Persistence and prevalence of pathogenic and extended-spectrum beta-lactamase-producing Escherichia coli in municipal wastewater treatment plant receiving slaughterhouse wastewater. Water Res 47:4719–29.

36. Blaak H, Hamidjaja RA, van Hoek AH, de Heer L, de Roda Husman AM, Schets FM. 2014. Detection of extended-spectrum beta-lactamase (ESBL)-producing Escherichia coli on flies at poultry farms. Appl Environ Microbiol 80:239–46.

37. Canton R, Akova M, Carmeli Y, Giske CG, Glupczynski Y, Gniadkowski M, Livermore DM, Miriagou V, Naas T, Rossolini GM, Samuelsen O, Seifert H, Woodford N, Nordmann P. 2012. Rapid evolution and spread of carbapenemases among Enterobacteriaceae in Europe. Clin Microbiol Infect 18:413–31.

38. Marti E, Huerta B, Rodriguez-Mozaz S, Barcelo D, Jofre J, Balcazar JL. 2014. Characterization of ciprofloxacin-resistant isolates from a wastewater treatment plant and its receiving river. Water Res 61:67–76.

39. Canton R, Novais A, Valverde A, Machado E, Peixe L, Baquero F, Coque TM. 2008. Prevalence and spread of extended-spectrum beta-lactamase-producing Enterobacteriaceae in Europe. Clin Microbiol Infect 14 Suppl 1:144–53.

40. Whitlock JE, Jones DT, Harwood VJ. 2002. Identification of the sources of fecal coliforms in an urban watershed using antibiotic resistance analysis. Water Res 36:4273–82.

41. Young KD, Thackston EL. 1999. Housing Density and Bacterial Loading in Urban Streams. J Environ Eng 125:1177–1180.

42. Kuhn I, Iversen A, Burman LG, Olsson-Liljequist B, Franklin A, Finn M, Aarestrup F, Seyfarth AM, Blanch AR, Vilanova X, Taylor H, Caplin J, Moreno MA, Dominguez L, Herrero IA, Mollby R. 2003. Comparison of enterococcal populations in animals, humans, and the environment--a European study. Int J Food Microbiol 88:133–45.

43. Reinthaler FF, Galler H, Feierl G, Haas D, Leitner E, Mascher F, Melkes A, Posch J, Pertschy B, Winter I, Himmel W, Marth E, Zarfel G. 2013. Resistance patterns of Escherichia coli isolated from sewage sludge in comparison with those isolated from human patients in 2000 and 2009. J Water Health 11:13–20.

44. Novo A, Manaia CM. 2010. Factors influencing antibiotic resistance burden in municipal wastewater treatment plants. Appl Microbiol Biotechnol 87:1157–66.

45. Institute CaLS. 2017. Performance Standards for Antimicrobial Susceptibility Testing: Twenty-seventh Informational Supplement M100-S27, CLSI, Wayne, PA, USA.

46. 2008 E-ESoCMaID. 2018. EUCAST: Clinical breakpoints. http://www.eucast.org/clinical_breakpoints/. Accessed

47. Lee K, Lim YS, Yong D, Yum JH, Chong Y. 2003. Evaluation of the Hodge test and the imipenem-EDTA double-disk synergy test for differentiating metallo-beta-lactamase-producing isolates of Pseudomonas spp. and Acinetobacter spp. J Clin Microbiol 41:4623–9.

48. Gupta G, Tak V, Mathur P. 2014. Detection of AmpC beta Lactamases in Gram-negative Bacteria. J Lab Physicians 6:1–6.

49. L. Biniossek SG, K. Xanthopoulou, E. Zander. M Kaase, H. Seifert, P. G. Higgins. 2016. Novel Multiplex PCR for detection of the most prevalent carbapenemase genes in Gram-negativebacteria within Germany. The 68th annual meeting of the German Society for Hygiene and Microbiology (DGHM).

50. Dallenne C, Da Costa A, Decre D, Favier C, Arlet G. 2010. Development of a set of multiplex PCR assays for the detection of genes encoding important beta-lactamases in Enterobacteriaceae. J Antimicrob Chemother 65:490–5.

51. Woodford N, Fagan EJ, Ellington MJ. 2006. Multiplex PCR for rapid detection of genes encoding CTX-M extended-spectrum (beta)-lactamases. J Antimicrob Chemother 57:154–5.

52. Rebelo AR MH, Cavaco L, Bortolaia V, Kjeldgaard JS, Hendriksen RS,. 2018. PCR for plasmid-mediated colistin resistance genes, mcr-1, mcr-2, mcr-3, mcr-4, mcr-5 and variants (multiplex), on DTU National Food Institute. https://www.eurl-ar.eu/CustomerData/Files/Folders/21-protocols/396_mcr-multiplex-pcr-protocol-v3-feb18.pdf. Accessed

53. Klindworth A, Pruesse E, Schweer T, Peplies J, Quast C, Horn M, Glockner FO. 2013. Evaluation of general 16S ribosomal RNA gene PCR primers for classical and next-generation sequencing-based diversity studies. Nucleic Acids Res 41:e1.

54. Dagert M, Ehrlich SD. 1979. Prolonged incubation in calcium chloride improves the competence of Escherichia coli cells. Gene 6:23–8.

55. Green R, Rogers EJ. 2013. Transformation of chemically competent E. coli. Methods Enzymol 529:329–36.

56. Johnson TJ, Nolan LK. 2009. Plasmid replicon typing, p 27-35. In Caugant DA (ed), Molecular Epidemiology of Microorganisms, 2009/06/13 ed, vol 551. Springer.

57. Martinez JL. 2009. Environmental pollution by antibiotics and by antibiotic resistance determinants. Environ Pollut 157:2893–902.

58. Control ECfDPa. 2015. SURVEILLANCE REPORT: Antimicrobial resistance surveillance in Europe 2014.

59. Ferreira da Silva M, Vaz-Moreira I, Gonzalez-Pajuelo M, Nunes OC, Manaia CM. 2007. Antimicrobial resistance patterns in Enterobacteriaceae isolated from an urban wastewater treatment plant. FEMS Microbiol Ecol 60:166–76.

60. Costa PMD, Vaz-Pires P, Bernardo F. 2008. Antimicrobial resistance in Escherichia coli isolated in inflow, effluent and sludge from municipal wastewater treatment plants. Urban Water Journal 4:275–281.

61. Costa EC, Arpini CM, Martins JDL. 2016. Antibiotic Sensitivity Profile of Enteric Bacteria Isolated from Beach Waters and Sewage from the Municipality of Vila Velha-ES, Brazil. Journal of Bacteriology & Parasitology 7:1–5.

62. Manaia CM, Novo A, Coelho B, Nunes OC. 2010. Ciprofloxacin Resistance in Domestic Wastewater Treatment Plants. Water, Air, and Soil Pollut 208:335–43.

63. Li D, Yu T, Zhang Y, Yang M, Li Z, Liu M, Qi R. 2010. Antibiotic Resistance Characteristics of Environmental Bacteria from an Oxytetracycline Production Wastewater Treatment Plant and the Receiving River^▿†^, p 3444–51, Appl Environ Microbiol, vol 76.

64. Du H, Chen L, Tang YW, Kreiswirth BN. 2016. Emergence of the mcr-1 colistin resistance gene in carbapenem-resistant Enterobacteriaceae. Lancet Infect Dis 16:287–8.

65. Liu YY, Wang Y, Walsh TR, Yi LX, Zhang R, Spencer J, Doi Y, Tian G, Dong B, Huang X, Yu LF, Gu D, Ren H, Chen X, Lv L, He D, Zhou H, Liang Z, Liu JH, Shen J. 2016. Emergence of plasmid-mediated colistin resistance mechanism MCR-1 in animals and human beings in China: a microbiological and molecular biological study. Lancet Infect Dis 16:161–8.

66. Malhotra-Kumar S, Xavier BB, Das AJ, Lammens C, Butaye P, Goossens H. 2016. Colistin resistance gene mcr-1 harboured on a multidrug resistant plasmid. Lancet Infect Dis 16:283–4.

67. Shen Z, Wang Y, Shen Y, Shen J, Wu C. 2016. Early emergence of mcr-1 in Escherichia coli from food-producing animals. Lancet Infect Dis 16:293.

68. Igwaran A, Iweriebor BC, Okoh AI. 2018. Molecular Characterization and Antimicrobial Resistance Pattern of Escherichia coli Recovered from Wastewater Treatment Plants in Eastern Cape South Africa, Int J Environ Res Public Health, vol 15.

69. Zurfuh K, Poirel L, Nordmann P, Nuesch-Inderbinen M, Hachler H, Stephan R. 2016. Occurrence of the Plasmid-Borne mcr-1 Colistin Resistance Gene in Extended-Spectrum-beta-Lactamase-Producing Enterobacteriaceae in River Water and Imported Vegetable Samples in Switzerland, p 2594–5, Antimicrob Agents Chemother, vol 60, United States.

70. Zhou HW, Zhang T, Ma JH, Fang Y, Wang HY, Huang ZX, Wang Y, Wu C, Chen GX. 2017. Occurrence of Plasmid- and Chromosome-Carried mcr-1 in Waterborne Enterobacteriaceae in China. Antimicrob Agents Chemother 61.

71. Drali R, Berrazeg M, Zidouni LL, Hamitouche F, Abbas AA, Deriet A, Mouffok F. 2018. Emergence of mcr-1 plasmid-mediated colistin-resistant Escherichia coli isolates from seawater. Sci Total Environ 642:90–94.

72. Tuo H, Yang Y, Tao X, Liu D, Li Y, Xie X, Li P, Gu J, Kong L, Xiang R, Lei C, Wang H, Zhang A. 2018. The Prevalence of Colistin Resistant Strains and Antibiotic Resistance Gene Profiles in Funan River, China. Front Microbiol 9:3094.

73. Munck C, Albertsen M, Telke A, Ellabaan M, Nielsen PH, Sommer MOA. 2015. Limited dissemination of the wastewater treatment plant core resistome. Nat Commun 6:8452.

74. White L, Hopkins KL, Meunier D, Perry CL, Pike R, Wilkinson P, Pickup RW, Cheesbrough J, Woodford N. 2016. Carbapenemase-producing Enterobacteriaceae in hospital wastewater: a reservoir that may be unrelated to clinical isolates. J Hosp Infect 93:145–51.

75. Harmon DE, Miranda OA, McCarley A, Eshaghian M, Carlson N, Ruiz C. 2019. Prevalence and characterization of carbapenem-resistant bacteria in water bodies in the Los Angeles-Southern California area. Microbiologyopen 8:e00692.

76. Lamba M, Gupta S, Shukla R, Graham DW, Sreekrishnan TR, Ahammad SZ. 2018. Carbapenem resistance exposures via wastewaters across New Delhi. Environ Int 119:302–308.

77. Zurfluh K, Bagutti C, Brodmann P, Alt M, Schulze J, Fanning S, Stephan R, Nuesch-Inderbinen M. 2017. Wastewater is a reservoir for clinically relevant carbapenemase- and 16s rRNA methylase-producing Enterobacteriaceae. Int J Antimicrob Agents 50:436–440.

78. Lefkowitz JR, Duran M. 2009. Changes in antibiotic resistance patterns of Escherichia coli during domestic wastewater treatment. Water Environ Res 81:878–85.

79. Garcia S, Wade B, Bauer C, Craig C, Nakaoka K, Lorowitz W. 2007. The effect of wastewater treatment on antibiotic resistance in Escherichia coli and Enterococcus sp. Water Environ Res 79:2387–95.

80. Gomi R, Matsuda T, Matsumura Y, Yamamoto M, Tanaka M, Ichiyama S, Yoneda M. 2017. Occurrence of Clinically Important Lineages, Including the Sequence Type 131 C1-M27 Subclone, among Extended-Spectrum-beta-Lactamase-Producing Escherichia coli in Wastewater. Antimicrob Agents Chemother 61.

81. Galvin S, Boyle F, Hickey P, Vellinga A, Morris D, Cormican M. 2010. Enumeration and characterization of antimicrobial-resistant Escherichia coli bacteria in effluent from municipal, hospital, and secondary treatment facility sources. Appl Environ Microbiol 76:4772–9.

82. Brechet C, Plantin J, Sauget M, Thouverez M, Talon D, Cholley P, Guyeux C, Hocquet D, Bertrand X. 2014. Wastewater treatment plants release large amounts of extended-spectrum beta-lactamase-producing Escherichia coli into the environment. Clin Infect Dis 58:1658–65.

83. Ojer-Usoz E, Gonzalez D, Garcia-Jalon I, Vitas AI. 2014. High dissemination of extended-spectrum beta-lactamase-producing Enterobacteriaceae in effluents from wastewater treatment plants. Water Res 56:37–47.

84. Caltagirone M, Nucleo E, Spalla M, Zara F, Novazzi F, Marchetti VM, Piazza A, Bitar I, De Cicco M, Paolucci S, Pilla G, Migliavacca R, Pagani L. 2017. Occurrence of Extended Spectrum beta-Lactamases, KPC-Type, and MCR-1.2-Producing Enterobacteriaceae from Wells, River Water, and Wastewater Treatment Plants in Oltrepo Pavese Area, Northern Italy. Front Microbiol 8:2232.

85. Chagas TP, Seki LM, Cury JC, Oliveira JA, Davila AM, Silva DM, Asensi MD. 2011. Multiresistance, beta-lactamase-encoding genes and bacterial diversity in hospital wastewater in Rio de Janeiro, Brazil. J Appl Microbiol 111:572–81.

86. Wu C, Wang Y, Shi X, Wang S, Ren H, Shen Z, Lin J. 2018. Rapid rise of the ESBL and mcr-1 genes in Escherichia coli of chicken origin in China, 2008-2014. Emerg Microbes Infect 7:30.

87. Marcade G, Deschamps C, Boyd A, Gautier V, Picard B, Branger C, Denamur E, Arlet G. 2009. Replicon typing of plasmids in Escherichia coli producing extended-spectrum beta-lactamases. J Antimicrob Chemother 63:67–71.

88. Sherley M, Gordon DM, Collignon PJ. 2003. Species differences in plasmid carriage in the Enterobacteriaceae. Plasmid 49:79–85.

89. Johnson TJ, Wannemuehler YM, Johnson SJ, Logue CM, White DG, Doetkott C, Nolan LK. 2007. Plasmid replicon typing of commensal and pathogenic Escherichia coli isolates. Appl Environ Microbiol 73:1976–83.

90. Boyd EF, Hill CW, Rich SM, Hartl DL. 1996. Mosaic structure of plasmids from natural populations of Escherichia coli. Genetics 143:1091–100.

91. Gniadkowski M. 2001. Evolution and epidemiology of extended-spectrum beta-lactamases (ESBLs) and ESBL-producing microorganisms. Clin Microbiol Infect 7:597–608.

